# An optimized three-laser 27-color spectral flow cytometry panel for multi-organ profiling in mice

**DOI:** 10.64898/2026.04.09.717400

**Authors:** Hyeonji Song, Yusik Lim, Jaechul Lim

## Abstract

While high-dimensional flow cytometry plays critical roles in resolving complex cellular networks, there remains a scarcity of comprehensive panels for the simultaneous profiling of diverse mouse cell types, primarily due to the inherent difficulty of multiplexing. To address this technical gap and resolve diverse cell populations in murine models, we designed a 27-color flow cytometry panel optimized for 3-laser spectral flow cytometers. This optimized panel enables broad and simultaneous detection of 16 distinct cell subsets from both lymphoid and myeloid lineages—including T cells, B cells, plasma cells, NK cells, innate lymphoid cells, dendritic cells, monocytes, macrophages, neutrophils, eosinophils, basophils, mast cells—along with non-immune cells, such as epithelial, endothelial, fibroblast, and neuronal cells. The panel has been successfully applied to various tissues, including spleen, thymus, bone marrow, peripheral blood, mesenteric lymph nodes, peritoneal lavage fluid, gut epithelium, and lamina propria. Applying this panel to a poly(I:C) model, we successfully tracked dynamic shifts in monocyte and neutrophil populations and identified a previously unrecognized, glucocorticoid-producing cell subset via reporter expression. This panel will facilitate high-dimensional immune profiling on standard 3-laser cytometers, providing a robust tool for dissecting cellular dynamics across diverse contexts.

## 1. Introduction

Mouse models are a cornerstone of modern immunology, offering unparalleled opportunities to investigate immune mechanisms at the cellular and organismal level (1). Unlike human studies that rely primarily on peripheral blood, murine systems allow direct analysis of immune responses within organs (2), thereby providing insight into tissue-resident and microenvironment-dependent immune regulation. Despite this advantage, most high-dimensional flow cytometry panels developed to date have concentrated on human samples, such as human PBMCs (3, 4), T cells, or antigen-presenting cells (5, 6). Consequently, there is a critical need for high-dimensional panels capable of capturing a broader range of murine immune cell populations. Notably, tissue-level immune analysis is particularly important owing to substantial heterogeneity in immune composition and activation states across lymphoid and non-lymphoid organs (7, 8). For example, mucosal tissues—such as the gut epithelium and lamina propria—are more complex than classical lymphoid organs, integrating inductive and tissue-resident compartments that balance tolerance and inflammation (9). Despite their significance, these regions have been underrepresented in prior panel designs due to complex tissue digestion protocols and the resulting poor cell viability and low yield (10). This need becomes even more apparent in systemic diseases such as sepsis, where dysregulated host responses extend across multiple organs and involve simultaneous alterations in diverse immune cell populations rather than a single dominant lineage (11). In these settings, immune dysregulation is accompanied by organ-specific inflammatory and metabolic changes, as well as inter-organ crosstalk, such that peripheral blood measurements alone may not fully reflect the tissue-level immune programs contributing to disease progression and organ dysfunction (12, 13). Accordingly, deep cell-level profiling across anatomically distinct tissues is essential for resolving how complex immune states are distributed across organs and for identifying coordinated or divergent cellular responses in systemic disease.

## 2. Materials and methods

### 2.1. Animals

Wild-type C57BL/6J (RRID: IMSR_JAX:000664) and homozygous *Cyp11b1-mScarlet* reporter mice (14) were maintained under SPF conditions with food and water ad libitum. Female mice aged 6–8 weeks were used in all experiments. Wild-type mice were used for panel optimization, including antibody titration and control setup, whereas *Cyp11b1-mScarlet* reporter mice were used for biological application experiments. All procedures were approved by the Institutional Animal Care and Use Committee (IACUC) of Seoul National University (SNU-240705-4-1) and conformed to institutional and national guidelines. Experiments were conducted and reported in accordance with ARRIVE guidelines.

### 2.2. Poly(I:C) treatment

*Cyp11b1-mScarlet* reporter mice were administered poly(I:C) (20 μg/g body weight; 2 mg/mL) via intraperitoneal injection (200 μL per mouse). Control *Cyp11b1-mScarlet* reporter mice received an equal volume of sterile saline. Tissues were collected 24 h after injection for flow cytometric analysis.

### 2.3. Instrument configuration

The panel was designed for a 3 Laser Cytek® Aurora spectral flow cytometer with the laser and detector specification displayed here (Table S1). Detector gains were applied according to the Cytek assay settings, with forward and side scatter voltages manually optimized to match the light scatter properties of the target cell populations.

### 2.4. Fluorochrome selection

Fluorochrome selection was guided by brightness index, spectral overlap characteristics, and instrument laser availability. Each fluorochrome was evaluated for spillover, spectral similarity, and signal resolution using Cytek’s Cloud Panel Builder (https://cloud.cytekbio.com) (Fig. S1). Fluorochrome assignment was adapted from the OMIP-069 (15) supplemental strategy but modified to ensure balanced spectral distribution across all lasers. Bright fluorochromes were assigned to low-abundance markers (RB744-CD117, PE-CD170), whereas dimmer fluorochromes were paired with highly expressed antigens (BV510-CD8, PerCP-Ly-6G). Cross-laser balance was further assessed using reference spectra to support accurate unmixing and minimize signal interference. Based on theoretical similarity and spreading error values provided by Cytek Cloud, fluorochrome combinations predicted to cause substantial spreading (>5.0) were selectively redistributed. In particular, assignments were optimized among APC/Alexa Fluor 647/Spark NIR685, RB744/PE-Cy7/RB705, and PerCP-Fire 806/BV785 to preserve resolution and minimize spreading error.

### 2.5. Reporter signal characterization

To incorporate the mScarlet reporter signal into the panel, the spectral profile of mScarlet was characterized using cells from *Cyp11b1-mScarlet* reporter mice. Its emission spectrum was compared with spectrally adjacent fluorochromes (PE, PE-eFluor 610, and PE-Fire 640) to evaluate potential overlap and confirm compatibility with the panel (Fig. S2).

### 2.6. Compensation

Compensation was performed using cell-based reference controls for each fluorochrome. Compensation beads were intentionally avoided, as previous studies and manufacturer documentation have reported subtle spectral discrepancies between antibodies bound to beads and those bound to cells (16, 17). All reference controls met the minimum criteria for both negative and positive event counts required by the SpectroFlo Unmixing Wizard, ensuring adequate signal separation and reliable reference spectra.

### 2.7. Antibody titration

Antibody titrations were performed in a 50 μL total staining volume per well. For each antibody, defined amounts were added according to the titration design, starting at 1 μg per test and followed by two-fold serial dilutions down to 0.0625 μg (i.e., 1 μg, 0.5 μg, 0.25 μg, 0.125 μg, and 0.0625 μg). Based on the unmixing results, additional lower concentrations were tested for fluorochromes that exhibited excessive spillover or signal saturation. For these antibodies, 3–5 additional dilution steps below 0.0625 μg were evaluated to identify the concentration that provided optimal separation between positive and negative populations while minimizing spectral spreading (Fig. S3 and Table S2).

All antibody titrations were initially performed using splenocytes, which represent a broad range of hematopoietic populations. However, several markers showed low or inconsistent expression in the spleen, resulting in insufficient positive event counts. For these markers, alternative tissues known to express the target antigen—such as bone marrow for CD31, CD56, CD161, and CD170 or intestinal epithelium for CD326—were used for titration under identical staining and acquisition conditions. This approach ensured reliable determination of optimal antibody concentrations across all fluorochromes, supporting the generation of robust reference spectra in subsequent unmixing steps.

### 2.8. Staining protocol

Staining was performed as follows:

1. Spin down and resuspend with 200 µL of ice-cold PBS.
2. Perform LIVE/DEAD violet (Thermo Fisher Scientific, Cat# L34964): 50 µL/sample (0.0031 µL LIVE/DEAD violet in 49.69 µL PBS), 30 min at RT in the dark, add 150 µL PBS and spin down.
3. Wash with 200 µL PBS per well, spin down as before.
4. Perform FC block (RRID: AB_2922498) (1:500): 50 µL/sample, 15 min at 4°C in the dark, add 150 µL FACS buffer and spin down.
5. Wash with 200 µL FACS buffer per well, spin down as before.
6. Perform Monocyte block (BioLegend, Cat# 426102): 50 µL/sample (2.5 µL Monocyte Blocker in 47.5 µL FACS buffer), 15 min at 4°C in the dark, add 150 µL FACS buffer and spin down.
7. Wash with 200 µL FACS buffer per well, spin down as before.
8. Prepare the staining master mix by combining optimal concentrations from single-color titrations according to the pre-calculated template. Use Brilliant Stain Buffer (RRID: AB_2869750) as the base when two or more BV fluorochromes are included.
9. Perform surface staining: 50 µL/sample, 30 min at 4°C in the dark, add 150 µL FACS buffer and spin down.
10. Wash with 200 µL FACS buffer per well, spin down as before.
11. Filter cells through 40 µm nylon mesh and transfer to FACS tube.

### 2.9. Autofluorescence extraction for spectral unmixing

To determine the optimal unmixing model, unstained controls from each tissue were analyzed using the Multiple AF Explorer function in SpectroFlo. Although unstained cells from different tissues showed broadly similar autofluorescence profiles, subtle tissue-specific differences in signal intensity were observed across some detectors. Unmixing without autofluorescence extraction left residual background signals, particularly in channels affecting population separation. Based on these comparisons, the Multiple AF Extraction (high AF) model was used for subsequent analyses to reduce tissue-derived background and improve spectral resolution (Fig. S4).

### 2.10. FMO controls for gate definition

Fluorescence minus one (FMO) controls were used for selected markers to support gate placement when positive and negative populations were not clearly separated after spectral unmixing. These controls were particularly useful for parameters affected by spectral spread and for defining gates for dim or infrequent populations with greater confidence (Fig. S5). Although FMOs were not generated for every marker, their targeted use helped improve gating consistency for selected populations.

### 2.11. Data analysis

All data were acquired on a 3-Laser Aurora system (Cytek Biosciences). Flow cytometry data were analyzed using FlowJo v10 (RRID: SCR_008520) and SpectroFlo v3.3.0 (RRID: SCR_025494). Dimensionality reduction was performed by UMAP (Uniform Manifold Approximation and Projection) using the FlowJo plugin, applied to gated cell populations from representative tissues.

## 3. Results

To address the limited availability of broadly applicable high-dimensional flow cytometry panels for diverse murine tissues, we developed a 27-color spectral flow cytometry panel for comprehensive cellular profiling using a standard 3-laser configuration (Fig. 1 and Table 1). Previously reported murine panels (18–28), have generally focused on lymphoid or myeloid subsets, or both only in limited settings, often within one or two organs and using four to five lasers. In contrast, our panel was designed to enable broad profiling across multiple murine tissues using a standard 3-laser spectral configuration.

**Fig. 1.**
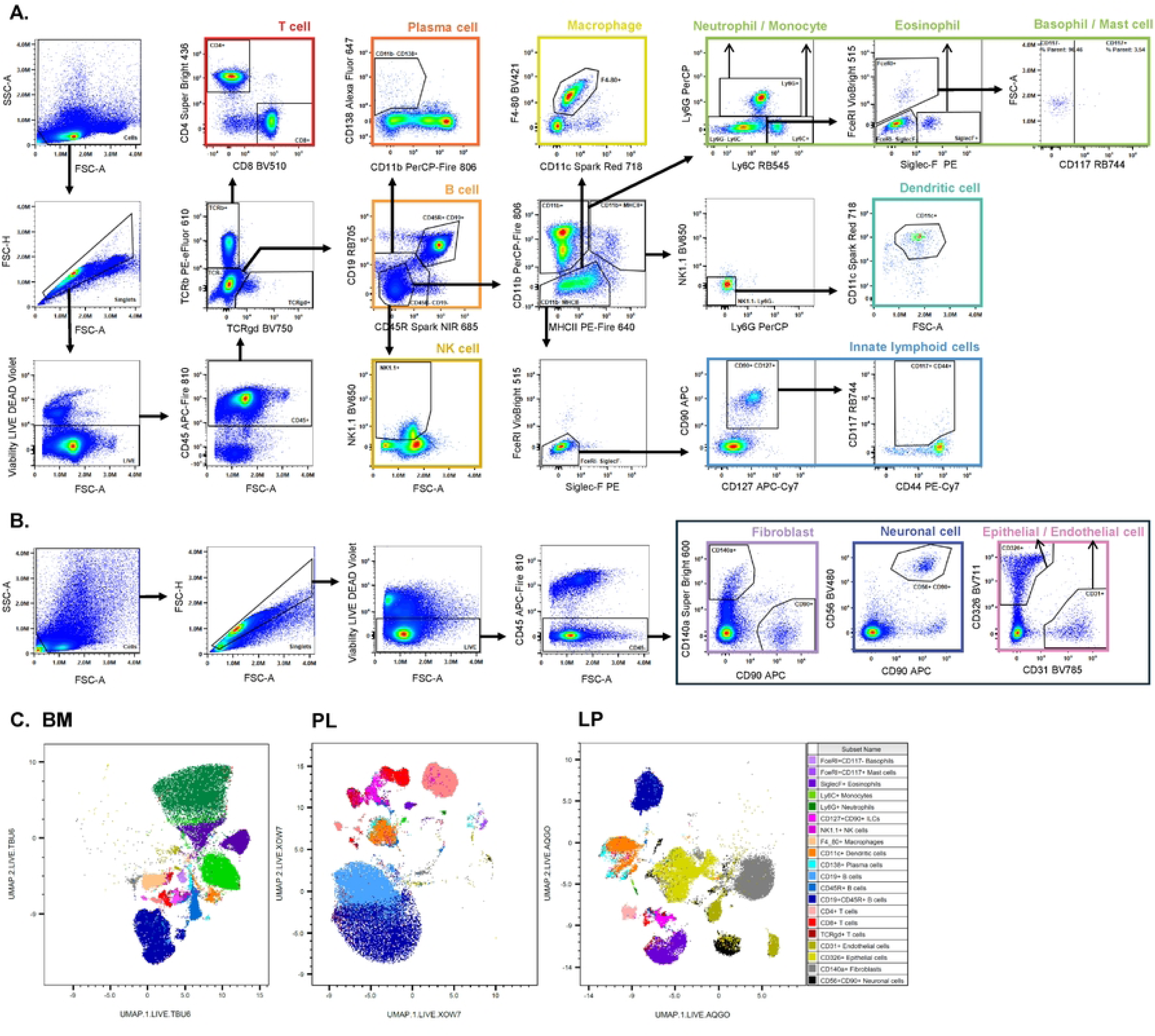
Gating Strategy. (A) Representative gating strategy for immune populations in the spleen. Populations were analyzed within the Cells/singlets/LIVE/CD45⁺ compartment and further resolved into major lymphoid and myeloid populations. (B) Representative gating strategy for non-immune populations in the small intestinal lamina propria. Populations were analyzed within the Cells/singlets/LIVE/CD45⁻ compartment to identify CD326⁺ epithelial cells, CD31⁺ endothelial cells, CD140a⁺ fibroblasts, and CD56⁺CD90⁺ neuronal cells. (C) UMAP visualization of gated cell populations from bone marrow (left, BM) and peritoneal lavage fluid (middle, PL), and lamina propria (right, LP). Cells were assigned to marker-defined populations based on the established gating strategy, and each color denotes a distinct annotated cell type shown in the legend.

**Table 1.**
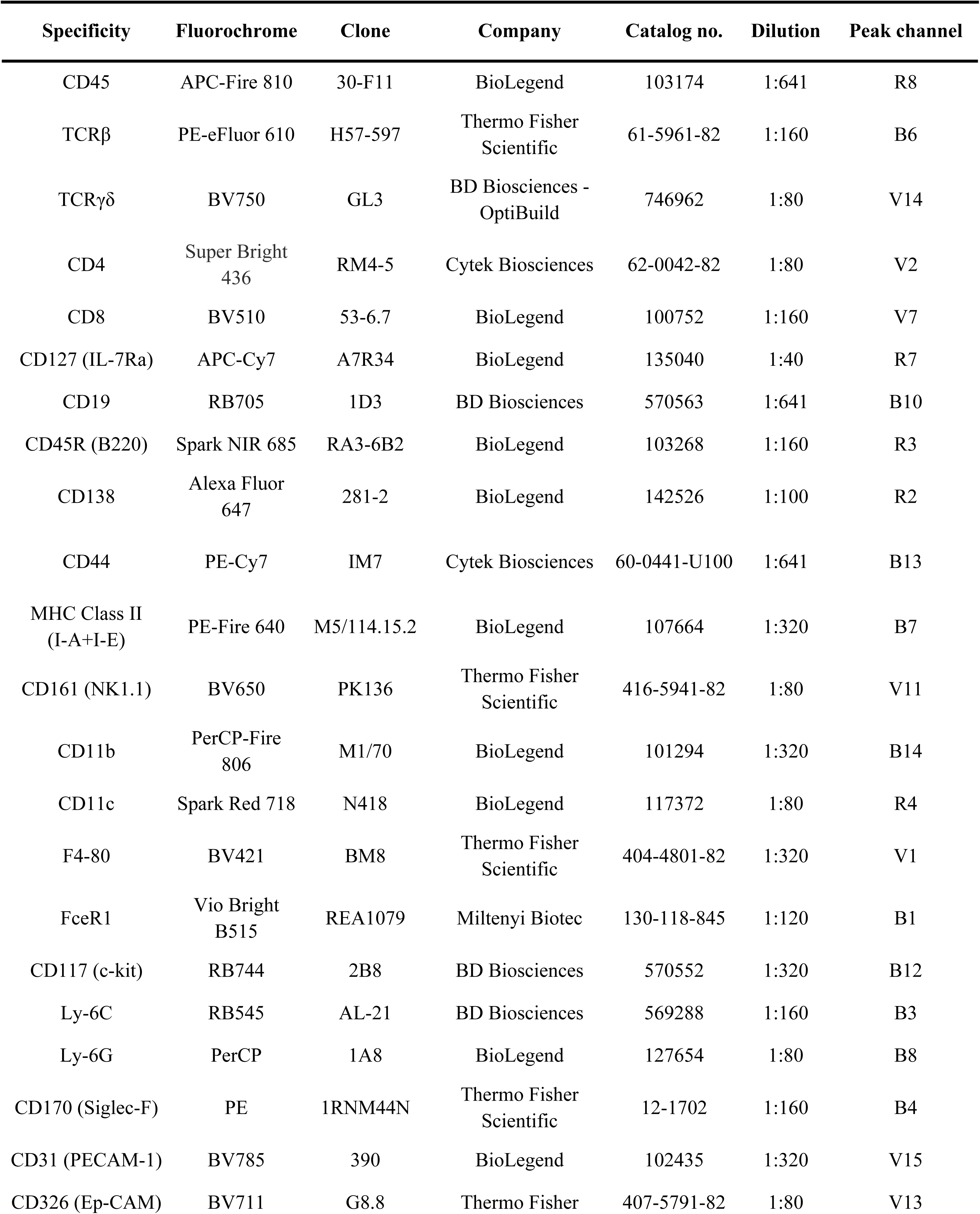
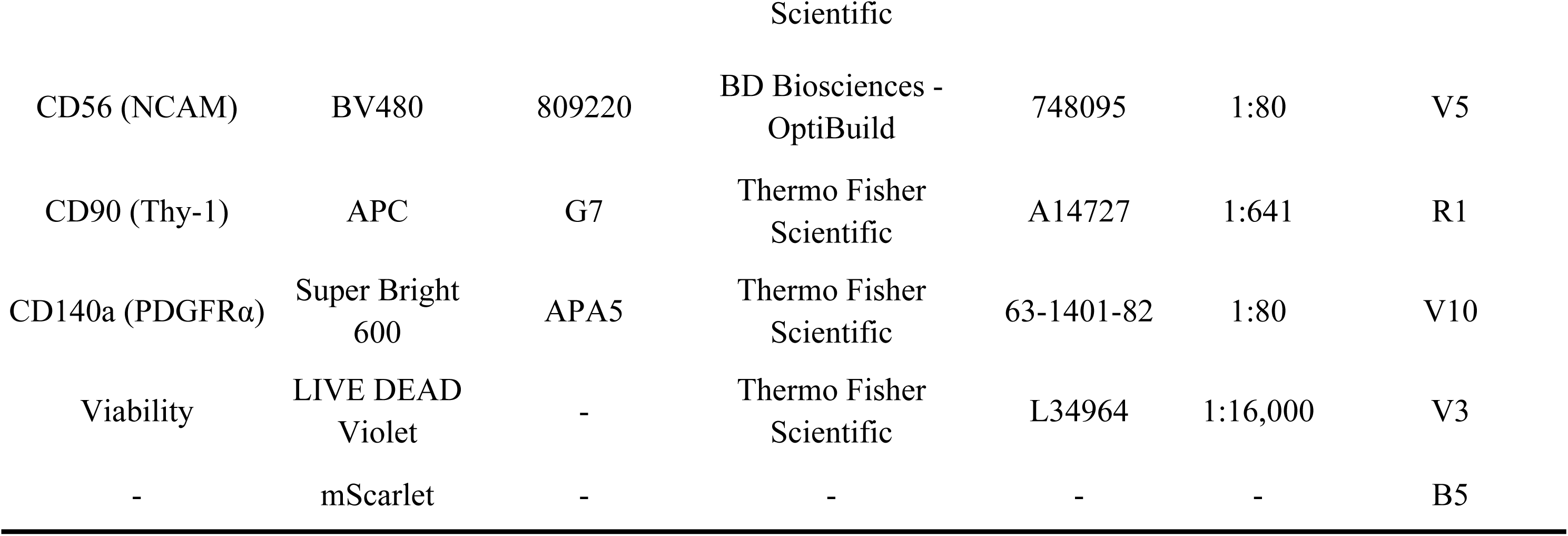
Antibodies used in the 27-color panel.

To construct the panel, markers were selected based on their expression patterns and immunological relevance, and fluorochromes were assigned with consideration of antigen density, signal resolution, and compatibility with spectral unmixing. Specifically, our panel is designed to simultaneously detect major lymphoid and myeloid lineages, using markers for T cells (CD4, CD8, TCRγδ), B cells (CD19, CD45R [B220]), plasma cell (CD138), NK cells (CD161 [NK1.1]), innate lymphoid cells (ILCs) (CD90 [Thy-1], CD127 [IL-7Ra], CD117 [c-Kit]), dendritic cells (CD11c), monocytes (Ly6C), macrophages (F4/80), neutrophils (Ly6G), eosinophils (CD170 [Siglec-F]), basophils, and mast cells (FcεRI, CD117 [c-Kit]), together with non-immune populations including endothelial cells (CD31 [PECAM-1]), epithelial cells (CD326 [EpCAM]), fibroblasts (CD140a [PDGFRα]), and neuronal cells (CD56 [NCAM], CD90 [Thy-1]). To guide panel architecture, markers were grouped into three hierarchical categories—primary, secondary, and tertiary—according to their roles in major lineage identification, subset discrimination, and phenotypic refinement (Table S3). Adapted from established immunophenotyping strategies, this framework provided a structured basis for panel design and interpretation of the cellular populations resolved by the panel while remaining compatible with spectral cytometry (6, 29).

Having established the panel architecture, we next optimized the panel to improve signal resolution and unmixing accuracy under full multi-color conditions. Specifically, antibody concentrations for selected markers were slightly adjusted, and single-stain reference files were refined prior to performance assessment. Because spectral overlap among fluorochromes inherently introduces spreading that can compromise signal resolution, we then evaluated the impact of spectral spread by comparing reference controls with multi-color spleen samples (Fig. S6). This comparison revealed that signal separation was noticeably reduced in the multi-color samples, particularly among fluorochromes with highly overlapping emission spectra, demonstrating the inherent resolution loss caused by cumulative spectral spreading in multi-color conditions. Unmixing performance was evaluated using NxN matrix plots across all fluorochrome combinations, and manual adjustments were applied when residual spillover was detected (Fig. S7). Across all fluorochrome combinations examined, correction values remained below 5%, indicating overall stable unmixing performance across tissues. The most prominent example was the APC–Alexa Fluor 647 pair, in which correction values reached up to 4.96% in epithelial samples and exceeded 2% in several tissues, including bone marrow, peripheral leukocytes, and lamina propria (Fig. S8). Similar but less pronounced patterns were observed for a subset of additional fluorochromes. In the case of CD90 APC, the recurrent deviation from Alexa Fluor 647 likely reflected a reference control that was not as bright as the corresponding sample signal, which is a known source of reduced unmixing accuracy. Despite these localized deviations, overall signal separation remained sufficient to reliably resolve the targeted cell populations across tissues. Therefore, this panel offers an integrated view of the murine cellular landscape within a single experiment.

Notably, it was initially developed as a 26-color antibody panel, but its configuration does not spectrally overlap with the endogenous mScarlet fluorescent reporter (14), allowing the inclusion of this fluorescent protein as an additional independent parameter, resulting in a 27-color panel. This design ensures broad compatibility across immune-focused mouse models while maintaining high dimensionality within a compact 3-laser setup. Detailed information on the markers and fluorophores is provided in Table 2. Taken together, these features provide a foundation for tissue-resolved cellular analysis in mice.

**Table 2.** Summary table for application of panel.

| Purpose | Profiling of murine immune and non-immune cells in multiple organs |
| --- | --- |
| Species | Mouse |
| Cell type | Single-cell suspensions or leukocytes isolated from murine spleen, thymus, mesenteric lymph nodes, bone marrow, peripheral blood, peritoneal lavage, gut epithelium, and lamina propria |
| Cross references | OMIP-31, OMIP-32, OMIP-57, OMIP-59, OMIP-61, OMIP-76, OMIP-86, OMIP-93, OMIP-95, OMIP-104, OMIP-105 |

To validate its utility, we next applied this panel to resolve major immune and non-immune cell populations across multiple murine tissues, including spleen, thymus, mesenteric lymph nodes, bone marrow, peripheral blood leukocytes, peritoneal lavage fluid, gut epithelium, and lamina propria, using a unified gating framework. Representative immune and non-immune gating strategies are shown (Figs. 1A and B). Following exclusion of debris, doublets, and dead cells, CD45⁺ immune cells and CD45⁻ non-immune cells were separated, allowing parallel analysis of lymphoid, myeloid, and stromal-associated compartments, respectively. Within the immune compartment, conventional αβ T cells were identified as TCRβ⁺TCRγδ⁻ cells and further resolved into CD4⁺ and CD8⁺ subsets, whereas γδ T cells were defined as TCRβ⁻TCRγδ⁺ cells. B cells were identified within the TCRβ⁻TCRγδ⁻ population based on CD19 and CD45R expression, including both CD19⁺CD45R⁺ double-positive and single-positive subsets. Within the remaining CD19⁻CD45R⁻ fraction, CD138⁺ plasma cells and NK1.1⁺ NK cells were identified. Among CD11b⁺MHC II⁺NK1.1⁻Ly6G⁻ cells, CD11c⁺ dendritic cells were detected, whereas within the CD11b⁻MHC II⁻ compartment, F4/80⁺ macrophages were identified. Within the CD11b⁻MHC II⁻FcεRI⁻SiglecF⁻ fraction, CD127⁺CD90⁺ cells corresponded to innate lymphoid cells (ILCs). Within the myeloid compartment, Ly6G⁺Ly6C⁻ cells were classified as neutrophils and Ly6G⁻Ly6C⁺ cells as monocytes. Among Ly6G⁻Ly6C⁻ cells, eosinophils were identified as FcεRI⁻SiglecF⁺ cells, while FcεRI⁺SiglecF⁻ cells were further separated into CD117⁺ mast cells and CD117⁻ basophils. Within the CD45⁻ compartment, epithelial cells were identified as CD326⁺ cells, endothelial cells as CD31⁺ cells, fibroblasts as CD140a⁺ cells, and neuronal-associated cells as CD56⁺CD90⁺ cells. This framework was further supported by UMAP visualization, in which cell populations defined by the established gating strategy formed distinct distributions in representative tissues such as bone marrow, peritoneal lavage fluid, and lamina propria (Fig. 1C). Thus, the panel provided a unified framework for broad cellular profiling across multiple murine tissues.

To evaluate the biological applicability, we applied the panel to an acute in vivo poly(I:C) model. Poly(I:C), a synthetic double-stranded RNA analog widely used as a viral mimic, is a well-established inducer of antiviral innate immune activation, including type I interferon–driven responses, and broader systemic immune perturbation (30). We therefore selected this model as a stringent test of whether our panel could capture coordinated immune remodeling across tissues in vivo. Using saline-treated and poly(I:C)-treated *Cyp11b1-mScarlet* reporter mice, we analyzed tissues 24 h after intraperitoneal injection. Broad immune cell changes were detected across multiple tissues, revealing substantial remodeling of systemic immune composition (Fig. 2). In the peripheral blood, peritoneal lavage fluid, and spleen, poly(I:C) exposure was associated with reduced CD4^+^ and CD8^+^ T cell frequencies, consistent with early redistribution of peripheral lymphoid compartments during acute antiviral-like inflammation (31, 32) (Fig. 2A). In parallel, Ly6G^+^ neutrophils and Ly6C^+^ monocytes increased across multiple tissues, in line with previous reports showing early mobilization and accumulation of innate myeloid populations following poly(I:C) exposure (33–35) (Fig. 2B). SiglecF^+^ eosinophils also showed a broad downward trend across tissues, with significant reductions in peripheral blood, peritoneal lavage fluid, spleen, and bone marrow (Fig. 2C). This pattern may reflect suppression or reprogramming of eosinophil-associated type 2 features under an antiviral/type 1-skewed inflammatory environment (36, 37). Together, these findings indicate that poly(I:C) drives a shift in immune composition away from a homeostatic lymphoid-dominant state toward a myeloid-biased inflammatory profile across tissues.

**Fig. 2.**
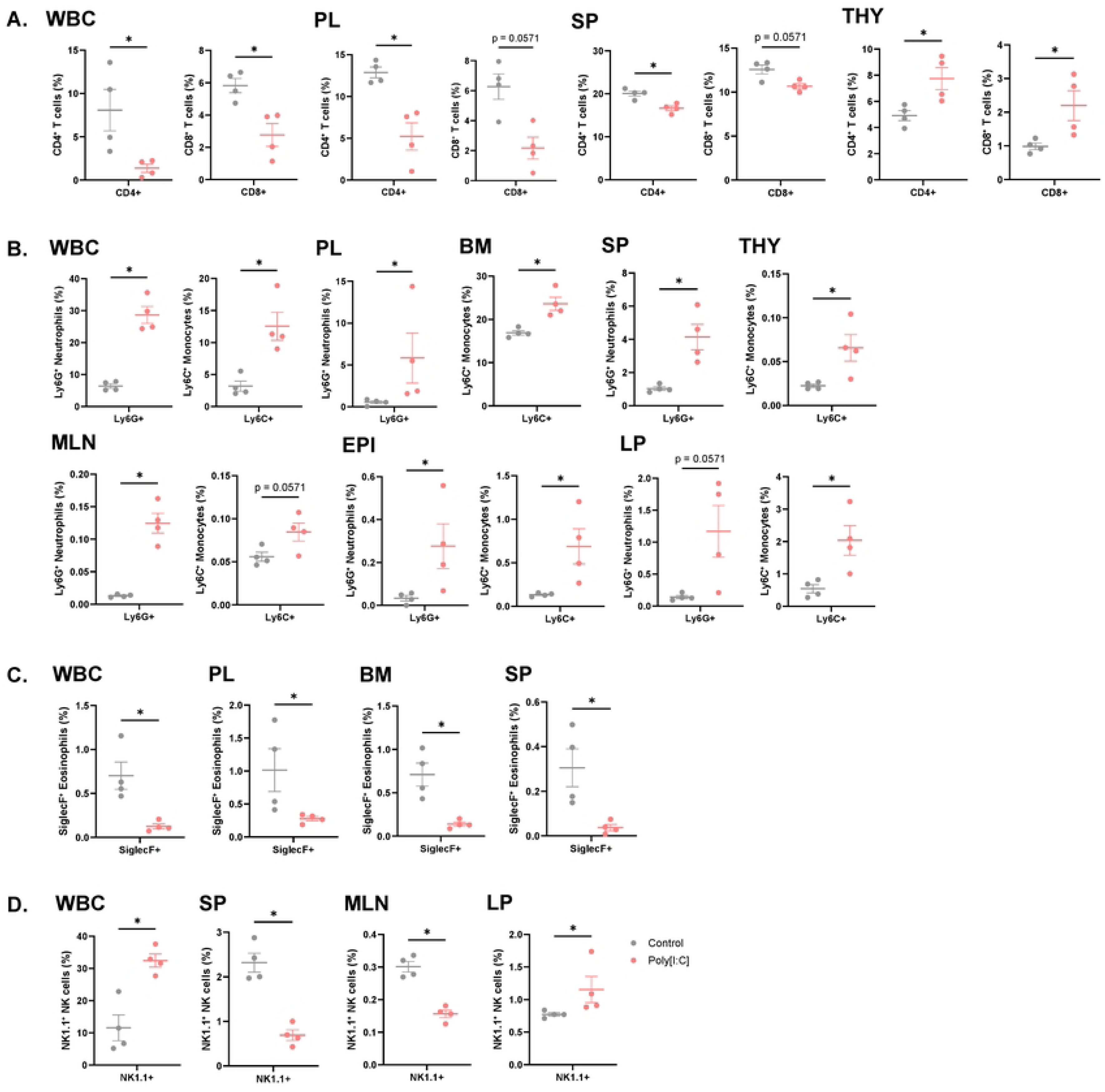
Cellular responses across tissues following poly(I:C) treatment. (A) Frequencies of CD4⁺ and CD8⁺ T cells across the indicated tissues following poly(I:C) treatment. Populations were analyzed within the CD45⁺/CD4⁺ and CD45⁺/CD8⁺ compartments. (B) Frequencies of Ly6G⁺ neutrophils and Ly6C⁺ monocytes across the indicated tissues following poly(I:C) treatment. Populations were analyzed within the CD45⁺/Ly6G⁺ and CD45⁺/Ly6C⁺ compartments. (C) Frequencies of SiglecF⁺ eosinophils across the indicated tissues following poly(I:C) treatment. Populations were analyzed within the CD45⁺/SiglecF⁺ compartment. (D) Frequencies of NK1.1⁺ NK cells across the indicated tissues following poly(I:C) treatment. Populations were analyzed within the CD45⁺/NK1.1⁺ compartment. WBC, white blood cells; PL, peritoneal lavage; BM, bone marrow; SP, spleen; THY, thymus; MLN, mesenteric lymph node; EPI, intestinal epithelium; LP, lamina propria. Each dot represents an individual mouse. Bars indicate mean ± SEM. Statistical significance was assessed using the Mann–Whitney test. *p < 0.05.

T cell responses were nevertheless compartment-specific. In contrast to the reduction in peripheral CD4^+^ and CD8^+^ T cells, thymic CD4^+^ and CD8^+^ T cell frequencies were maintained or increased, indicating that poly(I:C)-associated immune remodeling was not uniform across tissues (Fig. 2A). Because the thymus is the primary site of T cell development, where single-positive thymocytes undergo further maturation before export (38, 39), changes in thymic CD4^+^ and CD8^+^ frequencies may not directly reflect those observed in peripheral tissues. NK1.1^+^ NK cells also displayed tissue-dependent changes, with reduced frequencies in the spleen and the mesenteric lymph nodes but increased frequencies in the blood and the small-intestinal lamina propria, suggesting context-dependent redistribution or selective accumulation across compartments (Fig. 2D). Poly(I:C) is known to activate NK cells in vivo (40), although the tissue distribution of this response may vary with inflammatory context. These observations further illustrate the advantage of this panel, which enables direct comparison of the same cellular populations across multiple tissues within a single experiment. Collectively, the results demonstrate that the panel can resolve both common and tissue-specific features of systemic and intestinal immune remodeling in a biologically relevant antiviral-like inflammatory setting.

Beyond the broad immune changes detected across tissues, the panel also allowed endogenous mScarlet fluorescence to be analyzed in the same poly(I:C)-treated *Cyp11b1-mScarlet* reporter mice alongside antibody-based immune phenotyping, thereby enabling a full 27-color panel. Given that poly(I:C) elicits systemic inflammatory responses and is known to elevate circulating glucocorticoid levels (41, 42), this model provides a relevant context to examine whether Cyp11b1-associated reporter activity could be detected beyond the adrenal gland. Consistent with this, we found that mScarlet⁺CD4⁺ and mScarlet⁺F4/80⁺ are significantly increased by poly(I:C) challenge in the bone marrow and lamina propria of small intestine (Figs. 3A and B). These results indicate that T cells and macrophages may produce glucocorticoids during viral inflammation as previously described in tumors (43, 44). This increase in mScarlet⁺CD4⁺ T cells was therefore interpreted as a localized inflammatory response in bone marrow and lamina propria rather than a generalized T cell response. The F4/80⁺ pattern may reflect related changes along the monocyte–macrophage axis, in line with previous evidence that monocyte–macrophage lineage cells can participate in local glucocorticoid production (43). By contrast, interestingly, we discovered that Ly6C⁺ monocytes exhibiteda widespread increase in mScarlet expression across all tissues examined (Fig. 3C and Fig. S9). While previous studies have primarily focused on changes in monocyte abundance and activation state, our data further indicate that this response is accompanied by glucocorticoid production within monocytes. These results therefore indicate Ly6C⁺ monocytes as the most widespread and prominent glucocorticoid-producing immune population following poly(I:C) treatment, a finding that warrants further investigation. Collectively, these findings demonstrate that the panel is sensitive enough to resolve subtle endogenous reporter-associated shifts alongside broad immune phenotyping across tissues in vivo.

**Fig. 3.**
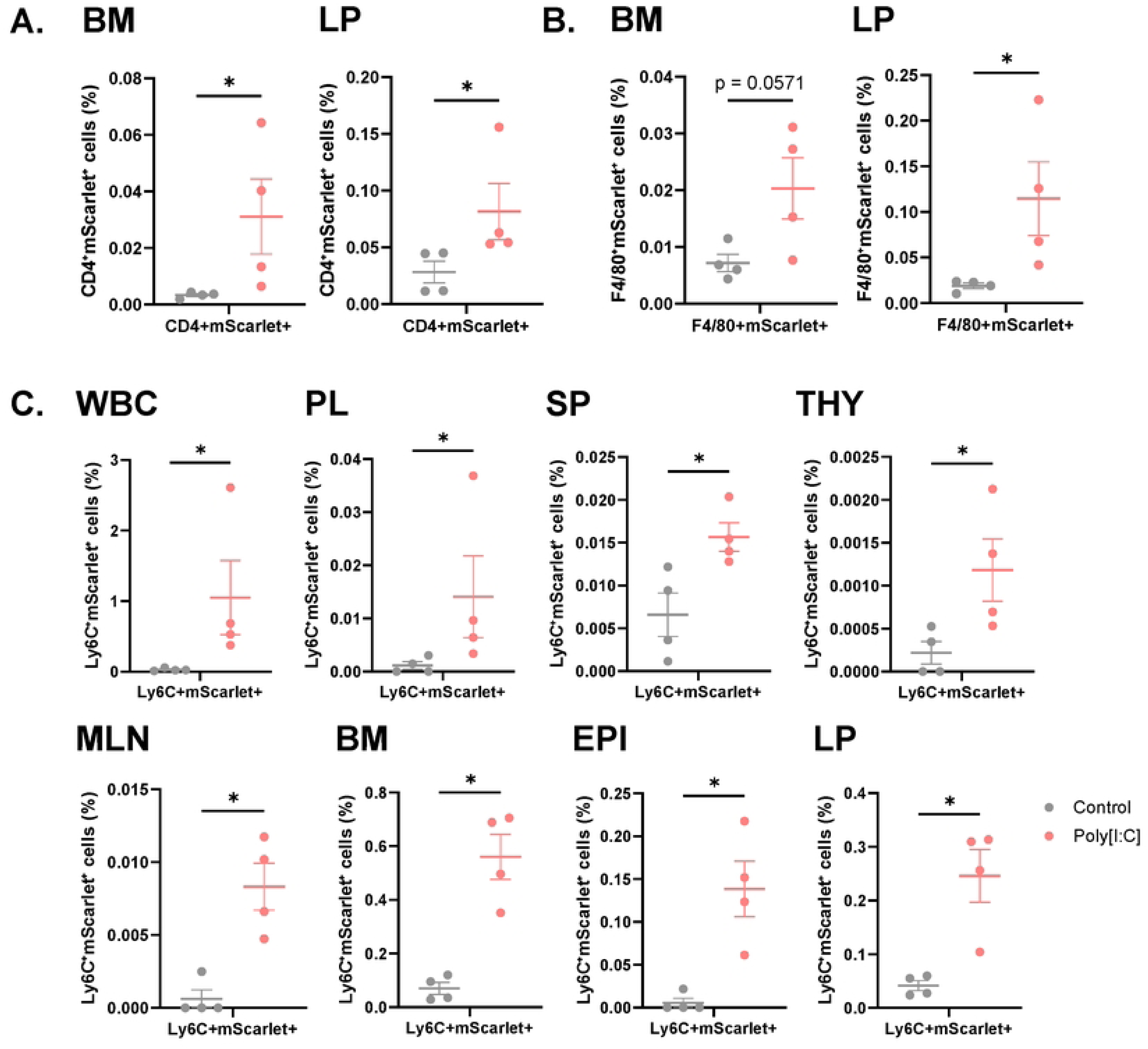
Compartment-specific mScarlet expression following poly(I:C) treatment. (A) Frequency of mScarlet⁺ cells in CD4⁺ immune cells in bone marrow and lamina propria following poly(I:C) treatment. Populations were analyzed within the CD45⁺/CD4⁺ compartment. (B) Frequency of mScarlet⁺ cells in F4/80⁺ immune cells in bone marrow and lamina propria following poly(I:C) treatment. Populations were analyzed within the CD45⁺/F4/80⁺ compartment. (C) Frequency of mScarlet⁺ cells in Ly6C⁺ immune cells across tissues following poly(I:C) treatment. Populations were analyzed within the CD45⁺/Ly6C⁺ compartment. WBC, white blood cells; PL, peritoneal lavage; SP, spleen; THY, thymus; MLN, mesenteric lymph node; BM, bone marrow; EPI, intestinal epithelium; LP, lamina propria. Each dot represents an individual mouse. Bars indicate mean ± SEM. Statistical significance was assessed using the Mann–Whitney test. *p < 0.05.

## 4. Discussion

In this study, we developed and optimized a 27-color full-spectrum flow cytometry panel for broad immunophenotyping in mice using a 3-laser platform. A key strength of this panel is that it enables simultaneous analysis of diverse immune populations together with selected non-immune cell types within a single workflow. This design is particularly advantageous in settings where tissue remodeling involves coordinated changes across multiple cellular compartments rather than isolated shifts within a single lineage. Accordingly, the panel is configured to capture major lymphoid and myeloid populations while also incorporating the Cyp11b1-mScarlet reporter, thereby extending conventional phenotyping to include glucocorticoid-associated cellular features within local tissue environments. This integrated design will be especially useful in models where systemic immune responses are accompanied by tissue-specific remodeling, including inflammatory, infectious, and stress-associated conditions. More broadly it provides a practical approach for examining how immune and structural compartments are jointly reorganized across multiple tissues or anatomical layers within the same mouse.

Notably, because the fluorochrome layout is spectrally optimized, this panel can be readily adapted for alternative targets—including intracellular or cell-type-specific markers— simply by substituting antibodies without major spectral re-optimization. By preserving this established spectral backbone, one can seamlessly transition from broad immunophenotyping to deep mechanistic studies. For instance, functional readouts such as transcription factors or cytokines can be integrated without disrupting the overall spectral balance. Relying on this pre-optimized configuration minimizes fluorochrome spreading errors and batch effects, providing a highly standardized and reproducible framework ideal for longitudinal tracking or multi-center collaborative studies. Moreover, the successful addition of an mScarlet reporter to the original 26-parameter panel exemplifies the expandability of our panel design. This indicates that other fluorescent proteins or dyes can be appended to accommodate diverse experimental designs.

Beyond this flexibility, these high-parameter capabilities are achieved using only a 3-laser full-spectrum configuration, enhancing practical accessibility and obviating the reliance on more instrument-intensive platforms. Importantly, despite this hardware constraint, the generated high-dimensional data retains the necessary depth and resolution to fully power advanced analytical pipelines. Consequently, researchers can readily apply sophisticated dimensionality reduction and clustering algorithms—such as t-SNE (t-distributed Stochastic Neighbor Embedding), UMAP, and trajectory inference—to dissect complex cellular networks using widely available cytometers.

In addition to its technical expandability and practical accessibility, the fundamental value of this high-dimensional panel lies in its capacity to capture complex biological and *in vivo* dynamics. We demonstrated its biological utility by uncovering the unexpected glucocorticoid-producing capacity of Ly6C⁺ monocytes during poly(I:C)-induced viral mimicry. While traditionally recognized primarily as potent drivers of acute inflammation, the identification of these myeloid cells as potential regulators in tissue-protective networks highlights the plasticity and dual-functional nature of the innate immune system. This biological insight has profound implications for a wide range of systemic inflammatory syndromes, such as sepsis, severe respiratory viral infections, and therapeutic cytokine release syndrome (CRS). Furthermore, this *in vivo* approach is equally vital for studying localized pathologies that exert systemic immunological consequences. For instance, in solid tumors or severe mucosal infections, localized tissue distress often triggers profound systemic immune alterations. As diverse immune populations are recruited to the disease microenvironment, the overall immune landscape undergoes significant reorganization. Furthermore, by incorporating representative non-immune lineages—such as epithelial or endothelial cells—our panel captures the spatial context of these immune infiltrates within the structural tissue niche. By enabling the high-resolution tracking of these population-level dynamics across multiple organs, our profiling strategy provides a critical tool for identifying systemic immune signatures in both systemic and localized disease conditions. Several limitations should also be noted. First, the panel is primarily designed for surface-marker–based profiling and does not include intracellular functional readouts such as cytokine production, transcription factor expression, or signaling activity. Thus, although it is effective for identifying changes in cellular composition, it is less suited for directly assessing functional states within each population. Second, although it captures a broad range of populations, some cell types are represented at a relatively general level and may require further refinement depending on the specific biological question. Additional marker substitution or panel adjustment may therefore be necessary for finer subclassification of closely related subsets. Finally, although the panel was optimized to minimize spectral interference, residual spreading was not completely eliminated in all channels. This likely reflects, in part, the inherent challenge of generating ideal reference controls for every marker under practical experimental conditions. In the present study, single-color controls were derived from tissue or cell populations exhibiting the strongest apparent signal within the sampled materials, which supported robust unmixing overall but may still leave room for further refinement.

## 5. Conclusions

In summary, the panel presented here enables parallel high-dimensional analysis of multiple murine cell populations across distinct tissue compartments using a compact 3-laser full-spectrum configuration. Its successful application in an acute poly(I:C) inflammatory model showed that it can resolve coordinated immune remodeling across tissues while simultaneously capturing subtle reporter-associated cellular states. These features make it a practical platform for studying complex immune and tissue-associated dynamics across diverse experimental settings.

## Authorship

Conceptualization: J.L., Formal Analysis: H.S., Y.L., Funding Acquisition: J.L., Investigation: H.S., Y.L., J.L., Methodology: H.S., Y.L., J.L., Visualization: H.S., Y.L., J.L., Writing – Original Draft Preparation: H.S., Y.L., J.L.

## Data availability

The data that support the findings of this study are openly available in Zenodo at https://doi.org/10.5281/zenodo.17873996, reference number 17873996.

## Acknowledgements and Funding information

This work was supported by the New Faculty Startup Fund (550-20230061), the Research Grant for Interdisciplinary Collaboration (550-20250081), the Research Institute for Veterinary Science, Seoul National University, and National Research Foundation (NRF), Ministry of Science, Information and Communication Technology (ICT) and Future Planning of Korea (RS-2021-NR057630, RS-2024-00335057 and RS-2024-00398705) (to J.L.).

## Supporting Online Material

An optimized three-laser 27-color spectral flow cytometry panel for multi-organ profiling in mice

---

**Fig. S1.**
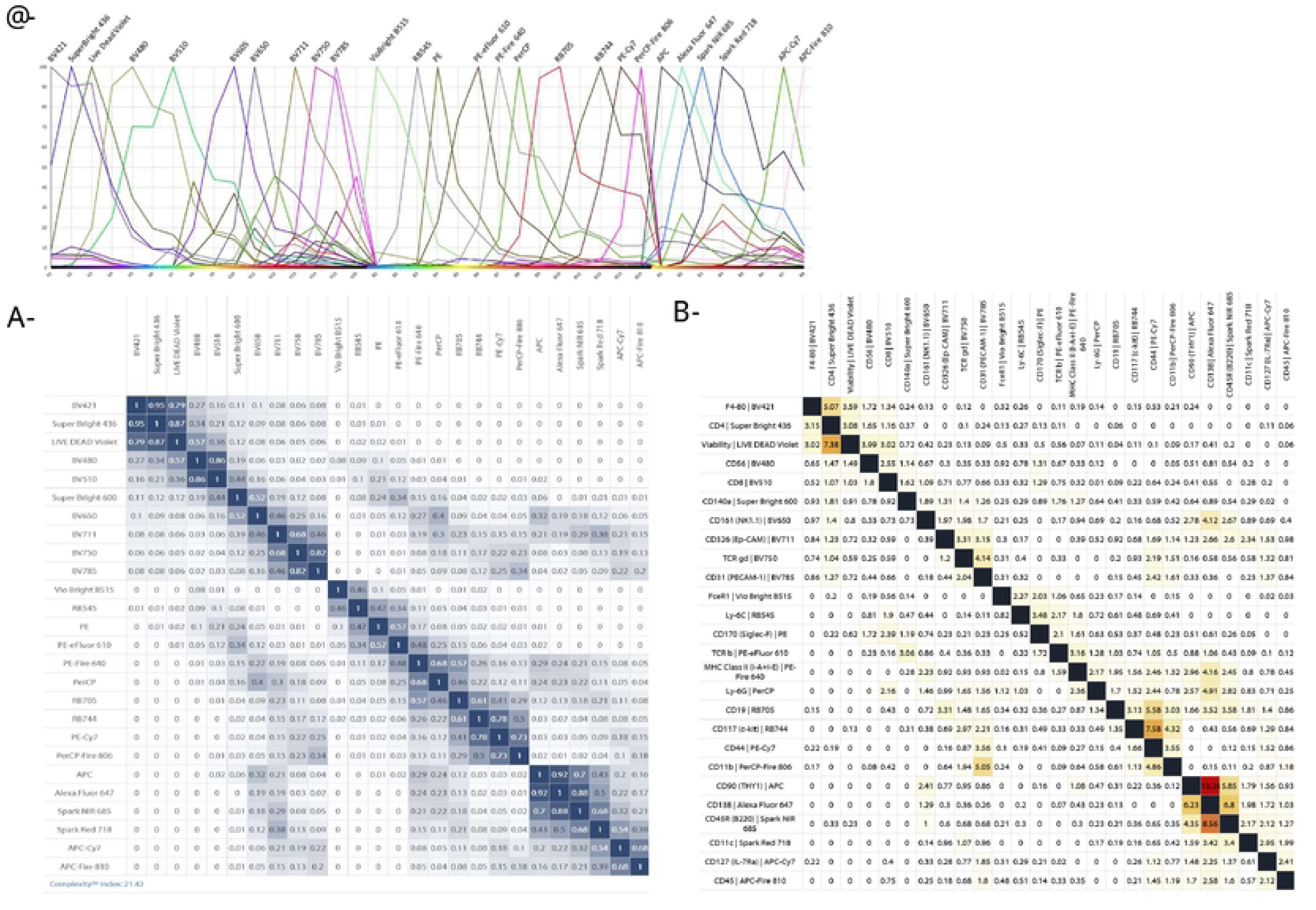
Spectral characteristics of fluorochromes included in the panel. (A) Emission spectra of all fluorochromes. (B) Similarity matrix displaying pairwise spectral overlap values between fluorochromes. The Similarity Index ranges from 0 (no spectral overlap) to 1 (identical spectra); values ≤ 0.98 indicate acceptable distinction between channels. The Complexity Index quantifies the cumulative spectral interference across all fluorochromes, providing an overall measure of panel complexity and predicting potential signal spread and autofluorescence impact. (C) Spillover spreading matrix (SSM) showing fluorescence spillover between detection channels, used to optimize panel design and minimize spectral overlap. Spectral characteristics were generated using Cytek’s Full Spectrum Viewer (https://spectrum.cytekbio.com).

**Fig. S2.**
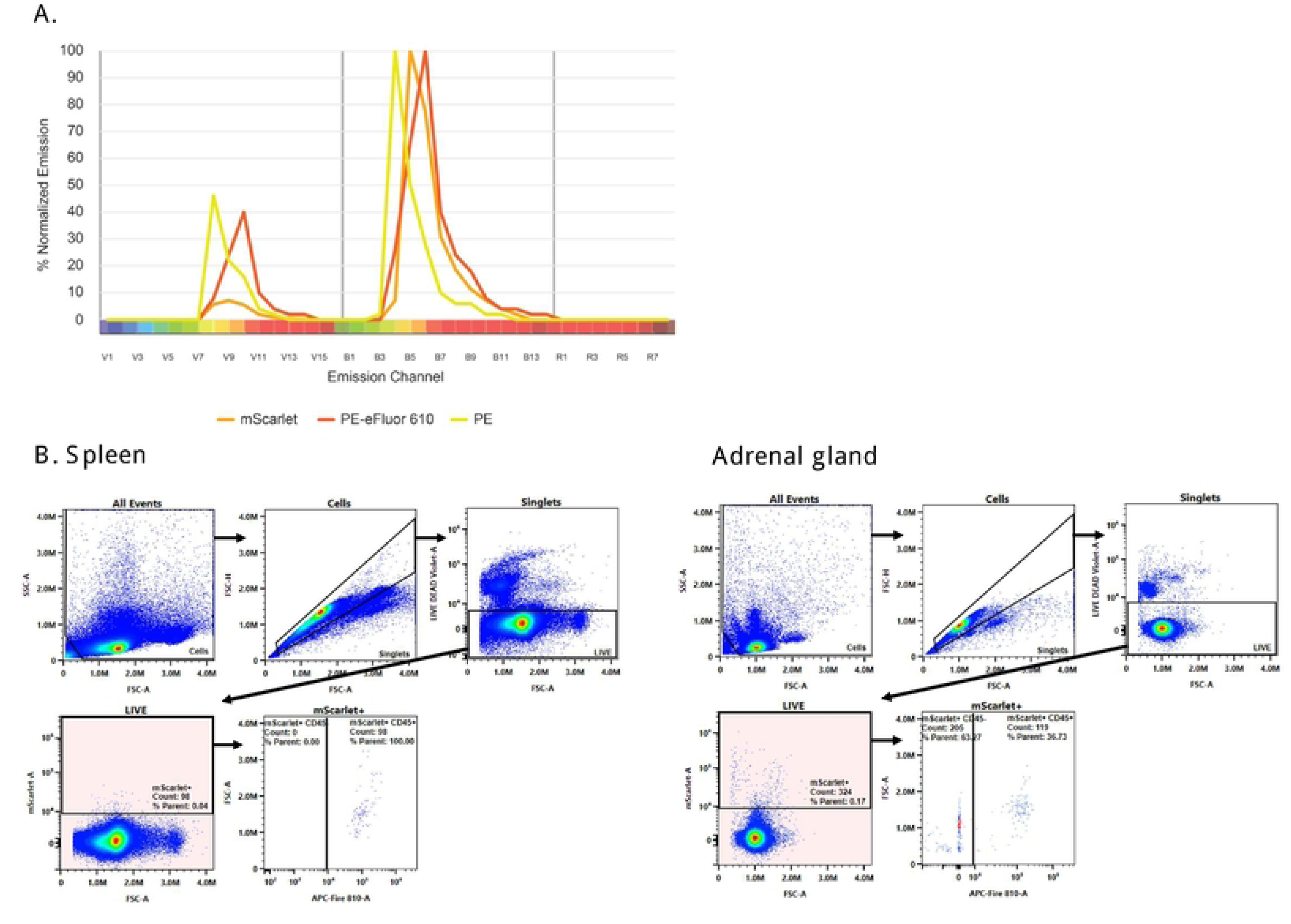
Spectral detection of mScarlet signal. (A) The emission spectrum of mScarlet is positioned between those of PE and PE-eFluor 610 (https://fluorofinder.com). (B) Gating strategy of spleen and adrenal gland cells from *Cyp11b1-mScarlet* reporter mice. Representative plots show sequential gating of Cells/Singlets/LIVE/mScarlet⁺ populations. In contrast to the spleen, mScarlet⁺ cells in the adrenal gland were detected mainly within the CD45⁻ fraction, consistent with reporter expression in non-immune parenchymal cells.

**Fig. S3.**
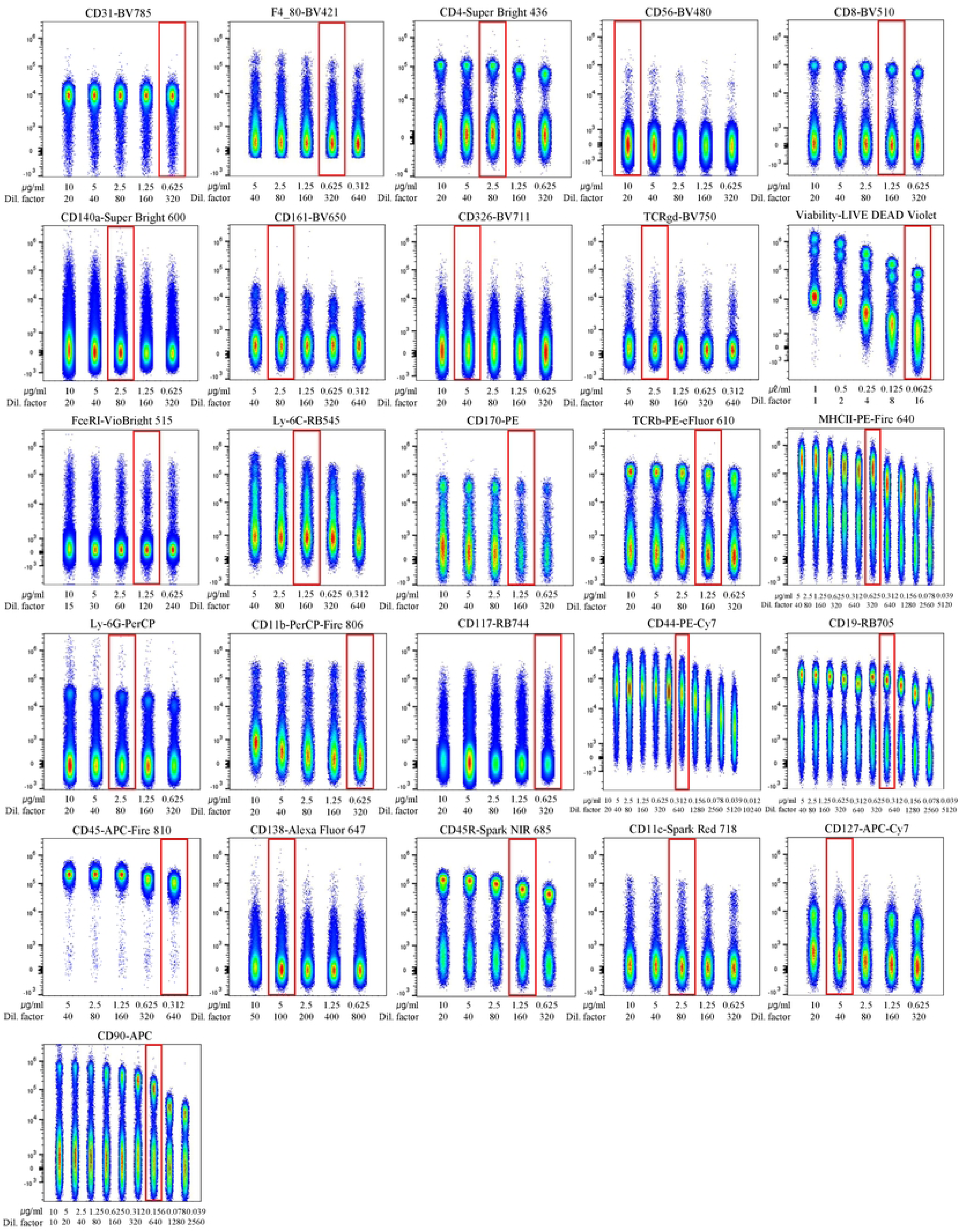
Titration of antibodies. All used antibodies were titrated to determine optimal concentrations. The x-axes of each plot denote the concentration of antibodies used in each titration condition. The y-axes show the fluorescence intensity of the given fluorochrome.

**Fig. S4.**
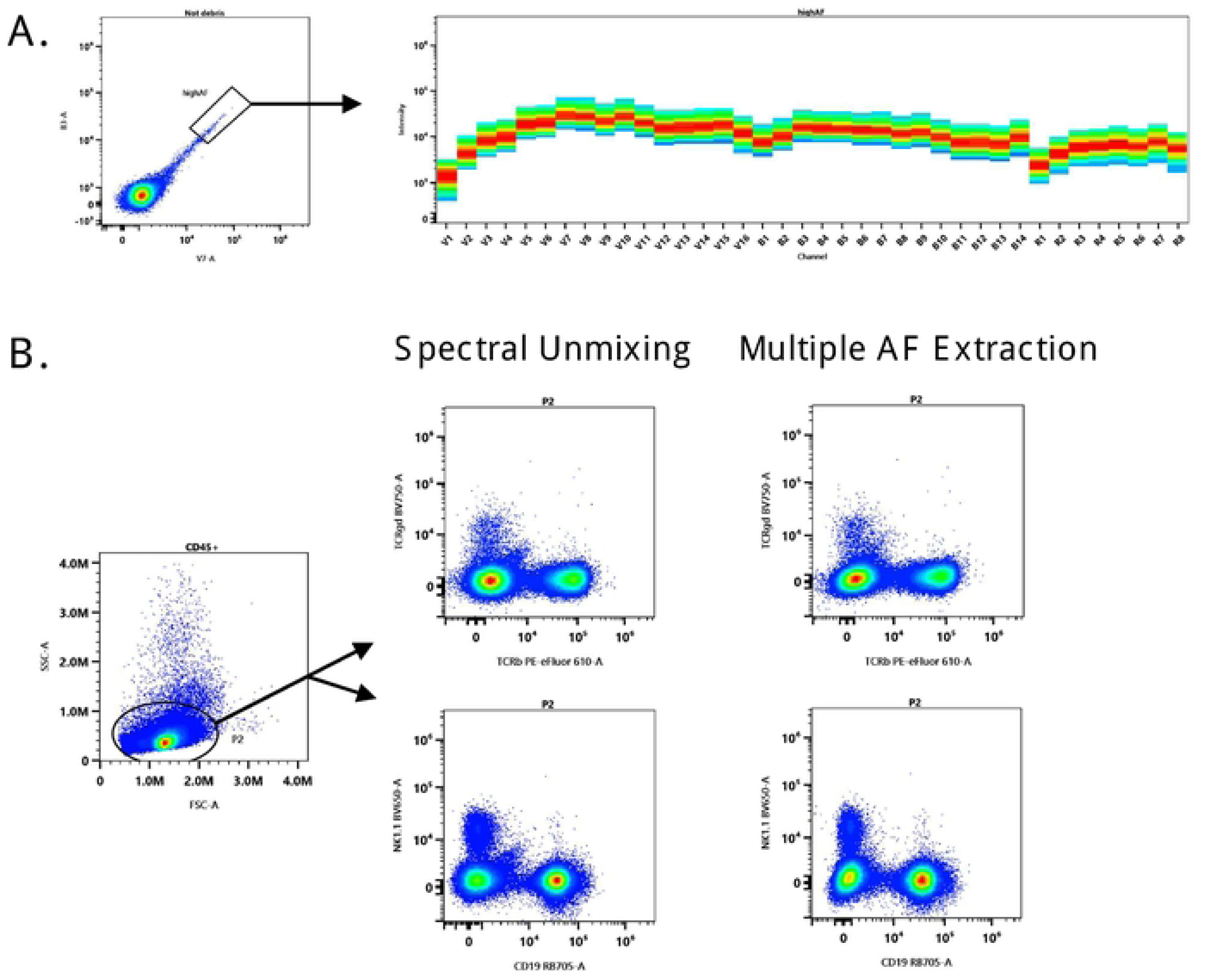
Unmixing model. (A) Autofluorescence assessment after removing doublets and debris, showing the gate defining the high-AF population and its corresponding spectral intensity profile. (B) Plots displayed after gating on Cells/Singlets/Live/CD45⁺, illustrating two representative regions where standard Spectral Unmixing and Multiple AF Extraction differ in background removal.

**Fig. S5.**
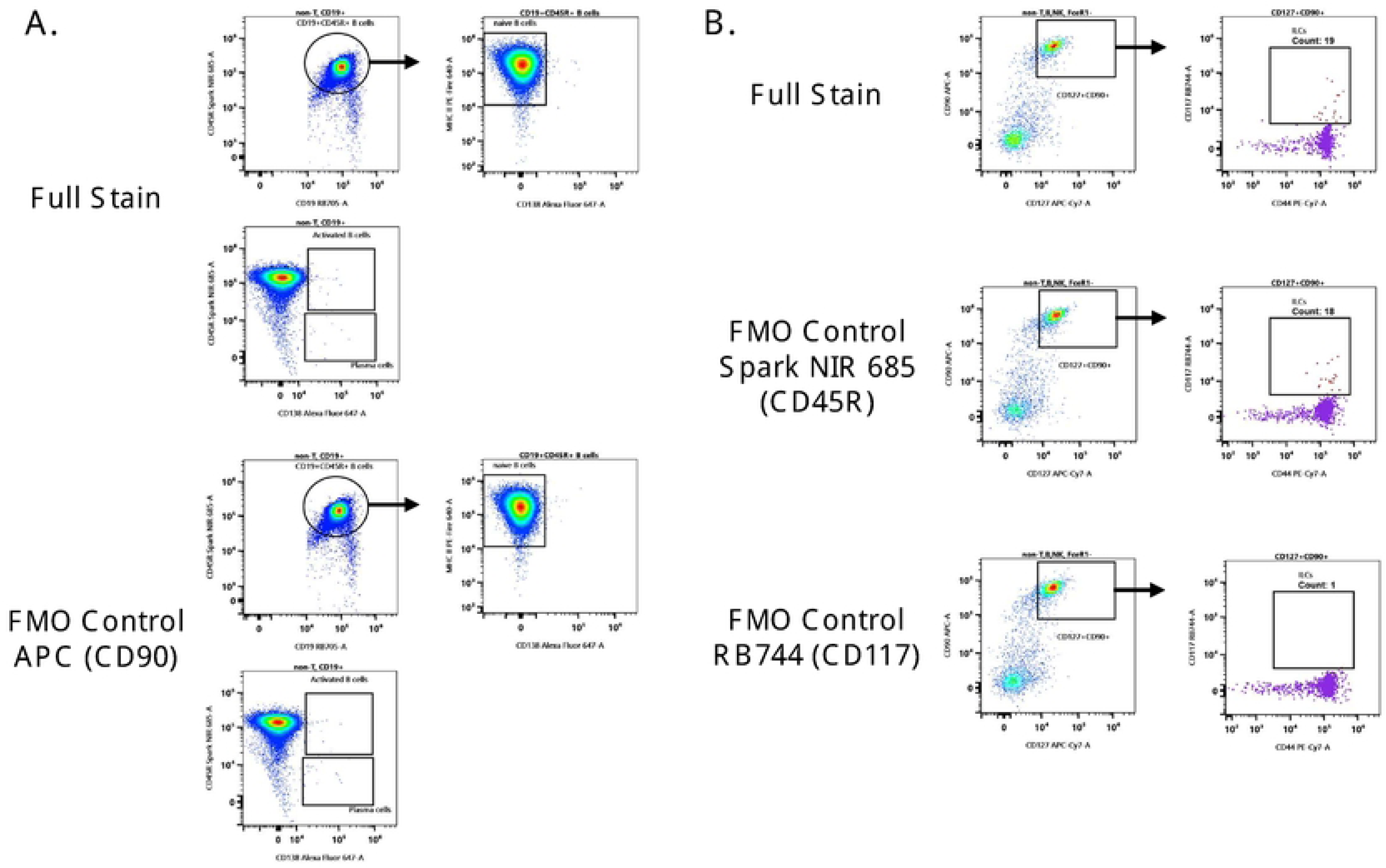
FMO Controls. (A) Representative full-stain and APC (CD90) fluorescence-minus-one (FMO) control plots showing gating of CD19⁺CD45R⁺ B cells within the TCRβ⁻TCRγδ⁻CD19⁺ compartment. Naïve B cells were identified from the CD19⁺CD45R⁺ population, whereas activated B cells and plasma cells were gated in parallel within the same parent gate. (B) Representative full-stain and fluorescence-minus-one (FMO) control plots for Spark NIR 685 (CD45R) and RB744 (CD117), showing identification of CD127⁺CD90⁺ cells within the TCRβ⁻TCRγδ⁻CD19⁻CD45R⁻NK1.1⁻FcεRI⁻ compartment. The right panels show the derived CD127⁺CD90⁺ population.

**Fig. S6.**
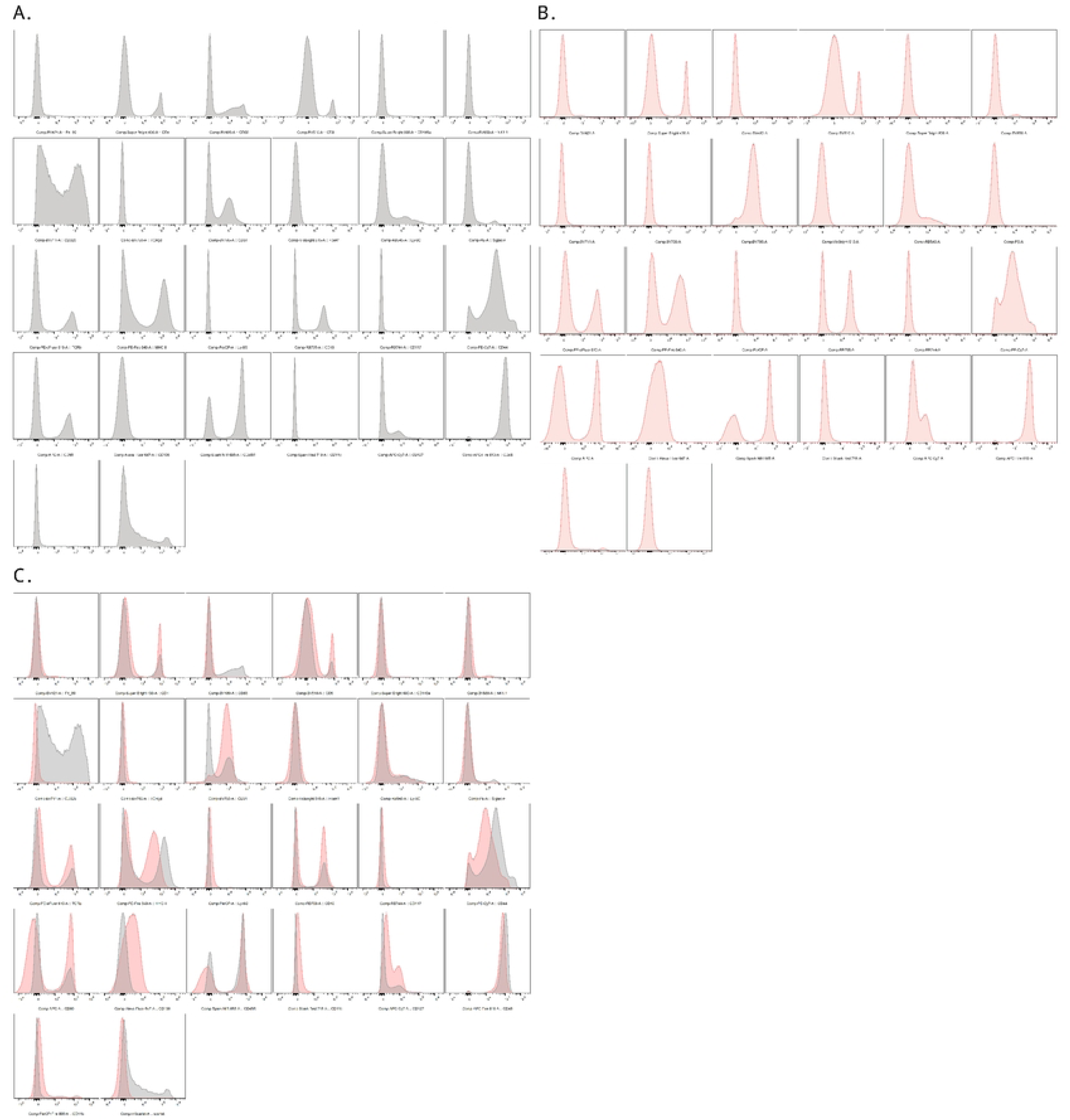
Signal resolution in multicolor samples and single-reference controls. (A) Histograms of the single-reference controls (RC) are shown for each of the 26 markers. (B) Corresponding histograms from multicolor (MC)–stained splenocytes gated on Cells/Singlets/Live are shown using the same markers and scaling. (C) Overlay of RC (gray) and MC (pink) histograms provides a direct marker-by-marker comparison of signal concordance and mismatch. Larger deviations for selected markers, such as BV711 (CD326), reflect the use of a reference control derived from a different tissue source.

**Fig. S7.**
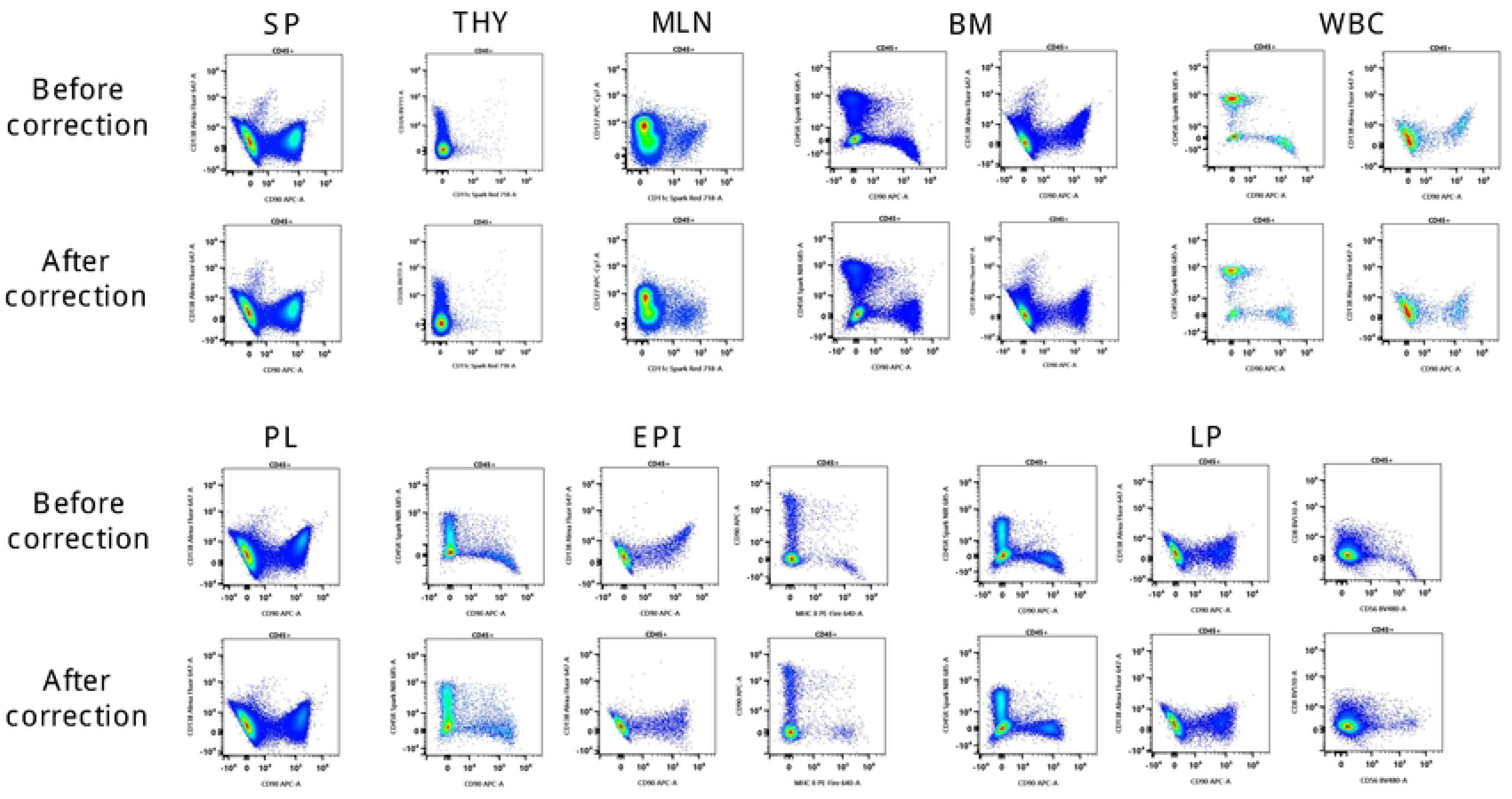
Compensation Correction. Compensation effects before and after correction are shown for events gated as Cells/Singlets/Live/CD45⁺ across multiple tissues. SP, spleen; THY, thymus; MLN, mesenteric lymph node; BM, bone marrow; PL, peritoneal lavage; WBC, white blood cells; EPI, intestinal epithelium; LP, lamina propria. Each pair of plots highlights a region where compensation changes the signal distribution.

**Fig. S8.**
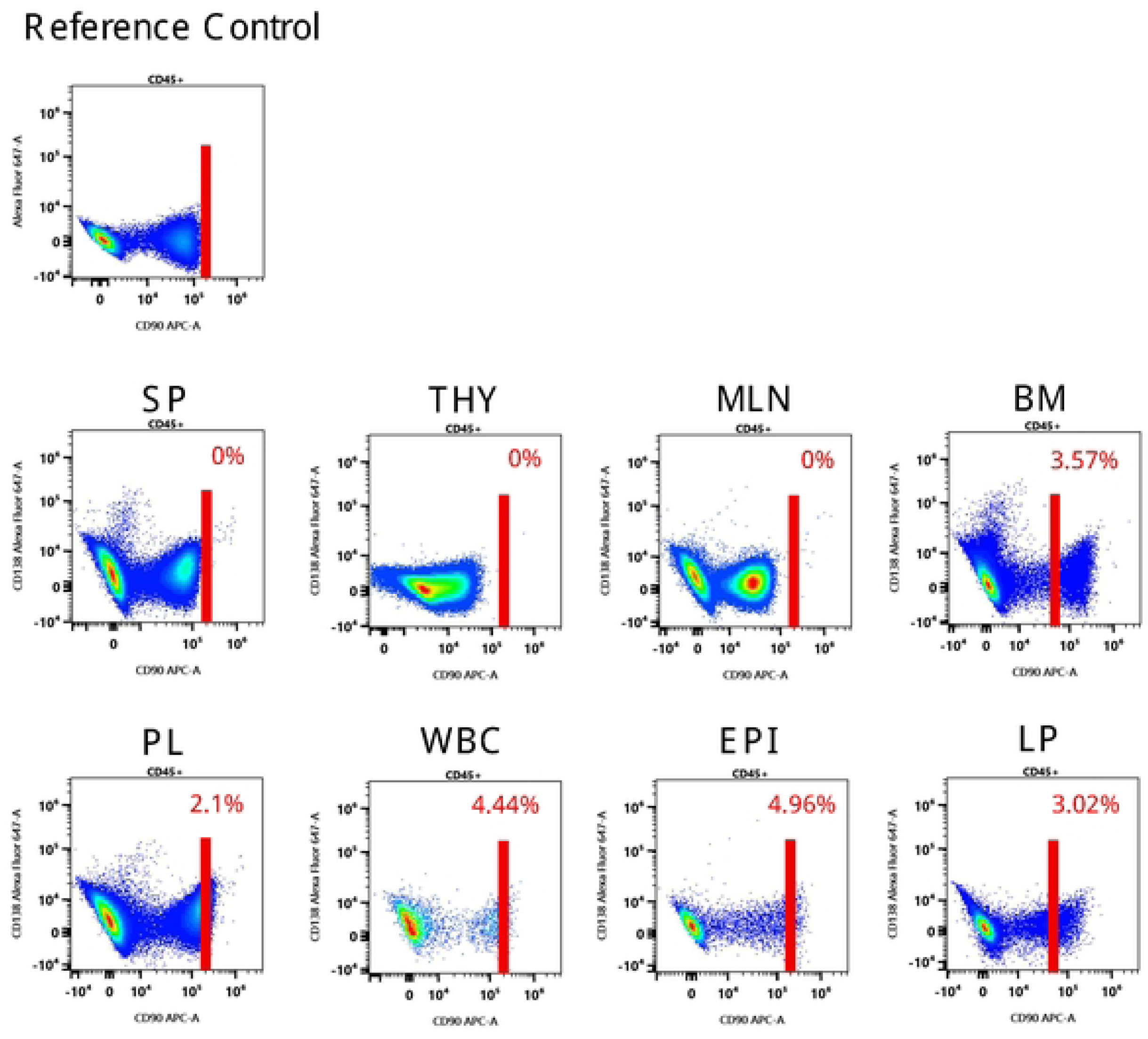
Comparison of marker MFI between multicolor samples and single-reference controls. Events gated on Cells/Singlets/Live/CD45⁺ are plotted on APC (CD90) and Alexa Fluor 647 (CD138) across multiple tissues. SP, spleen; THY, thymus; MLN, mesenteric lymph node; BM, bone marrow; PL, peritoneal lavage; WBC, white blood cells; EPI, intestinal epithelium; LP, lamina propria. A vertical cutoff (red line) derived from the reference control (RC) is applied to quantify the proportion of events exceeding this level in each multi-tissue control (MC), providing a visualization of pairwise compensation deviation for the APC–AF647 channel combination.

**Fig. S9.**
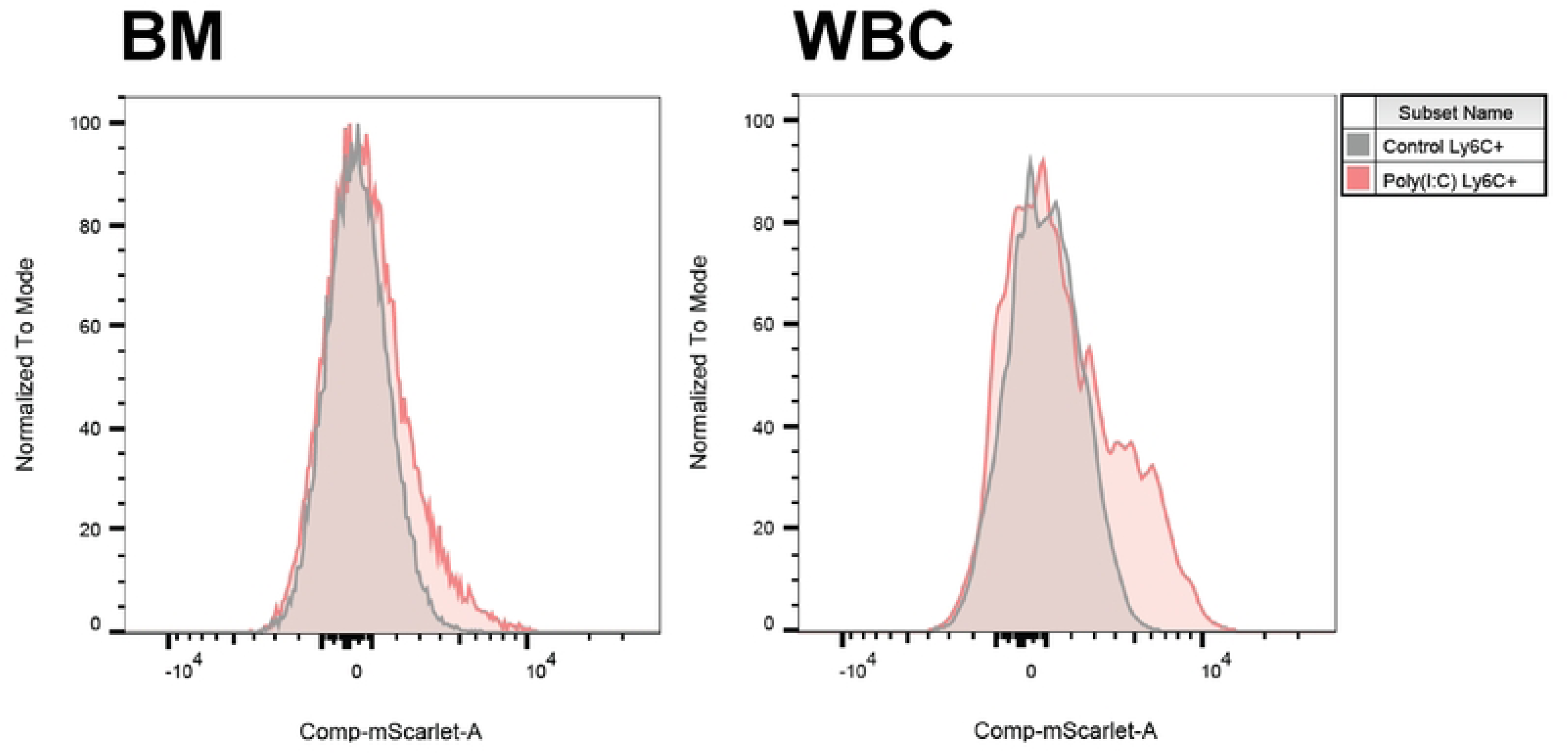
mScarlet expression in Ly6G⁺ cells following poly(I:C) treatment. Representative overlaid histograms showing mScarlet fluorescence intensity in CD45⁺/Ly6G⁺ cells in BM and WBC from control and poly(I:C)-treated mice. Control, gray; poly(I:C), pink. BM, bone marrow; WBC, white blood cells.

**Table S1.** Instrument configuration and setup for the Cytek 3-laser Aurora. Laser and detector configurations followed the standard settings provided by the manufacturer. The table summarizes the center wavelength and bandwidth (nm) of each detector channel, along with representative fluorochromes commonly assigned to each detector.

| Violet (405nm: 100mW) |  |  |  |
| --- | --- | --- | --- |
| Detector | Center Wavelength (nm) | Bandwidth (nm) | Fluorochrome |
| V1 | 428 | 15 | BV421 |
| V2 | 443 | 15 | SuperBright 436 |
| V3 | 458 | 15 | Live Dead Violet |
| V4 | 473 | 15 |  |
| V5 | 508 | 20 | BV480 |
| V6 | 525 | 17 |  |
| V7 | 542 | 17 | BV510 |
| V8 | 581 | 19 |  |
| V9 | 598 | 20 |  |
| V10 | 615 | 20 | SuperBright 600 |
| V11 | 664 | 27 | BV650 |
| V12 | 692 | 28 |  |
| V13 | 720 | 29 | BV711 |
| V14 | 750 | 30 | BV750 |
| V15 | 780 | 30 | BV785 |
| V16 | 812 | 34 |  |

| Blue (488nm: 50mW) |  |  |  |
| --- | --- | --- | --- |
| Detector | Center Wavelength (nm) | Bandwidth (nm) | Fluorochrome |
| B1 | 508 | 20 | VioBright B515 |
| B2 | 525 | 17 |  |
| B3 | 542 | 17 | RB545 |
| B4 | 581 | 19 | PE |
| B5 | 598 | 20 | mScarlet |
| B6 | 615 | 20 | PE-eFluor 610 |
| B7 | 661 | 17 | PE-Fire 640 |
| B8 | 679 | 18 | PerCP |
| B9 | 697 | 19 |  |
| B10 | 717 | 20 | RB705 |
| B11 | 738 | 21 |  |
| B12 | 760 | 23 | RB744 |
| B13 | 783 | 23 | PE-Cy7 |
| B14 | 812 | 24 | PerCP-Fire 806 |

| Red (638nm: 80mW) |  |  |  |
| --- | --- | --- | --- |
| Detector | Center Wavelength (nm) | Bandwidth (nm) | Fluorochrome |
| R1 | 661 | 17 | APC |
| R2 | 679 | 18 | Alexa Fluor 647 |
| R3 | 697 | 19 | Spark NIR 685 |
| R4 | 717 | 20 | Spark Red 718 |
| R5 | 738 | 21 |  |
| R6 | 760 | 23 |  |
| R7 | 783 | 23 | APC-Cy7 |
| R8 | 812 | 34 | APC-Fire 810 |

**Table S2.** Master mix for surface staining. Summary of dilution factors and volumes used to prepare the surface-staining master mix, with a total reaction volume of 50 µL.

| Fluorochrome (marker) | Dilution factor | Volume [μL]<br>in master mix |
| --- | --- | --- |
| APC (CD90, Thy-1) | 641 | 0.08 |
| APC-Cy7 (CD127, IL-7Rα) | 40 | 1.25 |
| APC-Fire 810 (CD45) | 641 | 0.08 |
| Alexa Fluor 647 (CD138) | 100 | 0.50 |
| Spark NIR 685 (CD45R, B220) | 160 | 0.31 |
| Spark Red 718 (CD11c) | 80 | 0.63 |
| BV421 (F4-80) | 320 | 0.16 |
| BV480 (CD56, NCAM) | 80 | 0.63 |
| BV510 (CD8) | 160 | 0.31 |
| BV650 (CD161, NK1.1) | 80 | 0.63 |
| BV711 (CD326, Ep-CAM) | 80 | 0.63 |
| BV750 (TCR$\gamma\delta$) | 80 | 0.63 |
| BV785 (CD31, PECAM-1) | 320 | 0.16 |
| Super Bright 436 (CD4) | 80 | 0.63 |
| Super Bright 600 (CD140a,<br>PDGFRα) | 80 | 0.63 |
| PE (CD170, Siglec-F) | 160 | 0.31 |
| PE-Cy7 (CD44) | 641 | 0.08 |
| PE-eFluor 610 (TCRβ) | 160 | 0.31 |
| PE-Fire 640 (MHC II) | 320 | 0.16 |
| PerCP (Ly-6G) | 80 | 0.63 |
| PerCP-Fire 806 (CD11b) | 320 | 0.16 |
| RB545 (Ly-6C) | 160 | 0.31 |
| RB705 (CD19) | 641 | 0.08 |
| RB744 (CD117, c-Kit) | 320 | 0.16 |
| Vio Bright B515 (FceRI) | 120 | 0.42 |

**Table S3.** Summary of markers and their classification. Summary of markers grouped into primary, secondary, and tertiary categories, with brief notes on their typical immunological roles.

| Marker | Classification | Notes |
| --- | --- | --- |
| CD45 | Primary | Strong leukocyte marker; distinct positive population. |
| TCRβ / TCR$\gamma\delta$ | Primary | Lineage-defining T cell marker; clear on/off expression. |
| CD4 / CD8 | Primary | T cell subset markers; discrete populations. |
| CD19 / CD45R (B220) | Primary | B cell lineage markers with clear gating separation. |
| Ly6G | Primary | Neutrophil marker; bright and unambiguous. |
| Ly6C | Secondary | Monocyte subset marker; continuum from high to low expression. |
| CD44 | Secondary | Activation and memory marker with graded expression levels. |
| CD161 (NK1.1) | Secondary | Strain-dependent NK marker; expression intensity varies. |
| CD11b / CD11c / F4/80 | Secondary | Myeloid markers with overlapping expression patterns. |
| FcεRI | Secondary | Mast and basophil marker; expression varies by tissue and activation state. |
| MHC II | Secondary | Antigen-presenting cell activation marker; variable intensity. |
| CD31 (PECAM-1) | Secondary | Endothelial/hematopoietic marker; moderate continuous expression. |
| CD170 (Siglec-F) | Secondary | Eosinophil marker; bright in some tissues; may behave as Primary in eosinophil-rich organs. |
| CD56 (NCAM) | Tertiary | Neuronal cell marker; low to moderate expression. |
| CD90 (Thy-1) | Tertiary | T cell, ILC, and neuronal marker; low or rare expression expected. |
| CD140a (PDGFRα) | Tertiary | Fibroblast and mesenchymal marker; moderate to variable expression. |
| CD127 (IL-7Rα) | Tertiary | ILC marker; expressed on innate lymphoid cells and variably on memory T cells. |
| CD138 | Tertiary | Plasma cell marker; limited tissue expression and frequency. |
| CD117 (c-Kit) | Tertiary | Mast cell and basophil lineage marker; low frequency, dim signal. |
| CD326 (EpCAM) | Tertiary | Epithelial cell marker; low or variable expression in mixed tissues. |

